# A Single Nucleotide Polymorphism assay sheds light on the extent and distribution of genetic diversity, population structure and functional basis of key traits in cultivated North American Cannabis

**DOI:** 10.1101/2020.02.16.951459

**Authors:** Philippe Henry, Surender Khatodia, Karan Kapoor, Britni Gonzales, Alexis Middleton, Kevin Hong, Aaron Hilyard, Steve Johnson, Davis Allen, Zachary Chester, Dan Jin, José Carlos Rodriguez Jule, Iain Wilson, Manu Gangola, Jason Broome, Deron Caplan, Dinesh Adhikary, Michael K. Deyholos, Michael Morgan, Oliver W. Hall, Brent Guppy, Cindy Orser

## Abstract

**Background:** The taxonomic classification of the Cannabis genus has been delineated through three main types: *sativa* (long and less branched plant with long and narrow leaves), *indica* (short but highly branched plant with broader leaves) and *ruderalis* (wild type with short stature, less branching and small thick leaves). While still under discussion, particularly whether the genus is polytypic or monotypic, this broad classification reflects putative geographical origin of each group and putative chemotypic and pharmacology.

**Methods:** Here we describe a thorough investigation of cannabis accessions using a set of 22 highly informative and polymorphic SNP markers associated with important traits such as cannabinoid and terpenoid expression as well as fibre and resin production. The assay offers insight into cannabis population structure, phylogenetic relationship, population genetics and correlation to secondary metabolite concentrations and demonstrate the utility of this assay for rapid, repeatable and cost-efficient genotyping of commercial and industrial cannabis accessions for use in product traceability, breeding programs, regulatory compliance and consumer education.

**Results:** The main outcomes are the identification of 5 clusters in the sample set available, including industrial hemp, resin hemp which likely underwent a bottleneck to stabilize CBDA accumulation (Type II & III). THC resin (type I) make up the other three clusters with terpinolene (colloquial “sativa” or “NLD”), myrcene/pinene and myrcene/limonene (colloquial “indica”, “BLD”), which also putatively harbour an active CBCAS.

**Conclusion:** The functional basis of key traits is also discussed as recently enabled by the NCBI Cannabis sativa Annotation Release 100, allowing for hypothesis testing with regards to secondary metabolite production as well as other key traits of importance for adaptable and compliant large-scale seed production under the new US Domestic Hemp Production Program.

## BACKGROUND

Cannabis, an annual and dioecious member of the family Cannabaceae, is an economically important genus providing protein- and oil-rich seeds, fibre biomass for industrial (construction, textile and paper) utilization, and a wide variety of secondary metabolites, predominantly terpenes and cannabinoids (Lynch et al., 2016; McPartland, 2018; Onofri and Mandolino, 2017). Cannabis produces over 150 types of terpenes and ∼100 different cannabinoids (Hanuš et al., 2016; Booth and Bohlman, 2019), however, its categorization into drug type and fibre type has historically been based mainly on a single cannabinoid, Δ^9^-tetrahydrocannabinol (THC). In this system, THC concentration (dry weight basis) >0.3% defines drug-type cultivars and ≤0.3% THC defines hemp cultivars (Dolgin 2019). This classification still prevails, including in the most recent USDA interim rules. Despite being grown and used for >6000 years in varying climates worldwide (Clarke and Merlin 2013), its evolution, taxonomic classification, and phylogenetic connections have been poorly understood. These deficiencies are due to limited genetic research, irregular breeding efforts, unorganized selection, *ex situ* conservation and government restrictions causing high heterozygosity in the cannabis genome (e.g. Rahn et al., 2016; McPartland 2018).

Taxonomic classification of the Cannabis genus has been delineated through three main types: *sativa* (long and less branched plant with long and narrow leaves), *indica* (short but highly branched plant with broader leaves) and *ruderalis* (wild type with short stature, less branching and small thick leaves). While still under discussion, particularly whether the genus is polytypic or monotypic, this broad classification reflects a putative geographical origin of each group (Clarke and Merlin 2017; Lynch et al., 2016, Schwabe et al., 2019). Consequently, there is no structured horticultural registration system available for Cannabis and cultivars or varieties, instead these are often awarded the epithet “strains”, which are likely the outcome of extensive hybridization of the original botanical descriptors (Henry, 2015).

Recent legalization of drug type cannabis for commercial production and recreational adult use in Canada, a number of US States and some other countries has brought about renewed scientific interest in developing a classification system for drug type cannabis. To that end, a particular focus has been placed on secondary metabolite expression with a clear separation based on CBD (cannabidiol):THC ratio, which is categorized into three classes: type I (<0.5), II (0.5-3.0) and III (>3.0) (Elzinga et al., 2015), A genetic basis for these types is determined by polymorphism at the CBDAS and THCAS genes on Chromosome 9 (Laverty et al., 2019). Double recessives at this locus would give rise to type IV (CBGA accumulators; de Meijer & Hammond, 2005). Type V plant would be cannabinoid-free chemotypes and may be the result of mutation in the upstream part of the cannabinoid synthase pathway (de Meijer et al., 2009). More recently the addition of terpenes as potential chemotaxonomic markers has emerged as a preferred model to cannabinoids alone (e.g. Lewis et al., 2018). Linking chemotype to genetic information has also enabled deeper insight into a novel consumer centric classification based on genetic markers associated with chemical expression (e.g. Orser and Henry, 2019). Recently, others have proposed targeted markers for the identification of fiber and resin Cannabis (e.g. Cascini et al., 2019; Hilyard et al., 2019) as well as molecular sexing tools to differentiate feminized from regular seed stock (Toth et al., 2020).

In addition to paving the way for informed classification, genetic information can also provide insights on the extent and distribution of genetic variability, population structure, phylogenetic relationships as well as providing the tools to shape a future breeding platform for cannabis with improved homozygosity and trait stability, and to identify clonal lines with identical multilocus genotypes. The latter may also be particularly useful in seed-to-sale tracking as it provides an irrefutable identity for each individual accession, possibly paving the way for cannabis variety registration and protection.

Here we describe a thorough investigation of cannabis accessions using a set of 22 highly informative and polymorphic SNP markers associated with important traits such as cannabinoid and terpenoid expression (Henry, 2017; Henry et al., 2018, Orser and Henry, 2019). We extend the scope of sampling to 681 accessions from licenced cultivators in Saskatchewan, Manitoba and British Columbia, Canada as well as Nevada, USA. We validated the use of these 22 SNP markers to assess population structure, phylogenetic relationship, population genetics and correlation to secondary metabolite concentrations and demonstrate the utility of this assay for rapid, repeatable and cost efficient genotyping of commercial and industrial cannabis accessions for use in product traceability, breeding programs, compliance and consumer education.

## METHODS

### Sample collection

Sample collection was undertaken to reflect the available diversity of cannabis germplasm available in North America, with samples from industrial hemp lines (type-III), resin hemp (type-II and type-III) and THC drug-type (type-I) Cannabis. Given the sensitivity of our genotyping approach, a small 2mm segment of leaf tissue was sufficient to yield adequate DNA for downstream genotyping.

### DNA Isolation procedure

Prior to performing the DNA extraction protocol, and in order to obtain high molecular weight DNA, plant tissue samples were allowed to air dry for 24-48hrs at room temperature and in the presence of silica desiccant. Plant tissue was homogenised in a 1.5ml Eppendorf tube with a reusable pestle. Homogenised material was then treated following the Sbeadex® plant mini kit protocol (LGC Biosearch Technologies, Beverley, MA) following the manufacturer’s instructions. Briefly, after the addition of 90µL Lysis buffer PN, samples were incubated at 65 °C for >10 minutes. The samples were then centrifuged at 2500 x g for 10 minutes to pellet the debris. 50µL of the supernatant in this tube, referred to as the lysate was then transferred to another 1.5ml tube with 120µL Binding buffer PN and 10µL Sbeadex® particle suspension and incubated at room temperature for 4 minutes. The tube was then brought into contact with a magnet for about a minute until the magnetic particles form a pellet. The supernatant was then discarded and the pellet was then subjected to three consecutive wash steps. The washed beads were then eluted with 70µL Elution buffer PN and incubated at 55 °C for 3 minutes prior to bringing the tubes in contact with the magnet. 50µL of the eluate was then transferred to a new tube which contain high purity plant DNA.

### Endpoint PCR genotyping using custom KASP assays

Twenty-two optimized assay mixes, each specific to single nucleotide polymorphisms (SNP) previously identified as associated with phylogeny and chemotypic expression were screened in the sample set (Henry 2015, 2017; Henry et al., 2018). These assays consist of two competitive, allele-specific forward primers and one common reverse primer (KASP; LGC Biosearch Technologies, Beverley, MA). Each forward primer incorporates an additional tail sequence that corresponds with one of two universal FRET (fluorescent resonance energy transfer) cassettes present in the KASP Master mix which contains the two FRET cassettes (FAM and HEX), ROX™ passive reference dye, Taq polymerase, free nucleotides and MgCl_2_ in an optimised buffer solution.

The genotypes were generated using an Eco RT (Illumina, San Diego, CA), a CFX 96 (Biorad, Hercules, CA) and an Intelliqube array tape platform (LGC Biosearch Technologies, Beverley, MA) with multiple blind replicates across platforms to ensure cross system repeatability. Genotypes were called using the Kluster Caller software and manually verified using the SNPviewer software (LGC Biosearch Technologies, Beverley, MA).

### Functional basis of 22 SNP

Given the recent release of the 10 chromosome map of the cannabis genome (Grassa, 2018; Laverty et al., 2019), metabolomic and proteomic insight (Jenkins and Orsburn, 2019a,b) as well as a fully annotated version of the cannabis genome resulting from the completion of the NCBI Cannabis sativa Annotation Release 100 (Jenkins and Orsburn, 2019c), we set out to characterise the functional basis of the SNPs used in the study. The previously designed targets developed using Cansat 3 (von Bakel et al., 2011) were subjected to a BLASTn search (Altschul et al., 1990) constrained to the taxa Cannabis using the NCBI online interface (https://blast.ncbi.nlm.nih.gov) accessed October 31, 2019. The location of the 10 chromosome map as well as the putative functional gene in which the 22 SNP are found were recorded.

### Statistical Analyses of genotypic data

Multilocus genotypes were formatted as a table (comma separated file) of genotypes with individuals as rows and markers as columns. As the total dataset of 681 plant DNA samples contained some missing data, we culled all missing data out and undertook the following analyses on 420 samples with complete genotype information across all markers. Metadata, including individual and population names, were separated from the genotype data and imported into the flexible statistical environment of R (R development core team 2018) requiring the following packages, *ape* (Paradis & Schliep, 2018), *pegas* (Paradis, 2010), *poppr* (Kamvar et al., 2014), *adegenet* (Jombart, 2008) and *hierfstat* (Goudet and Jombart, 2015).

Briefly, the *read.loci* function was used to import the allelic data into the R environment as a data frame which was then converted to a *genind* object using the *df2genind* command. Individual and population (variety identity) were also incorporated into the *genind* object to allow for population level calculations to shed light on the stability of claimed variety names and to assess the level of genetic diversity within and between these hypothesized groups. Clonal lines were identified using *mlg* and *mlg.id* functions, which determines the number and identity of mutilocus genotypes. Basic population genetics metrics, particularly expected heterozygosity were calculated for each population and individuals using the *poppr* function.

To shed light on the underlying relationships between our diverse sample set, a dissimilarity matrix or Hamming distance between multilocus genotypes was calculated using the *bitwise.dist* function and was visualized using a phylogenetic tree using the *nj* function. Principal component analyses were undertaken to provide an independent line of evidence of the genetic affinities between accessions using the *dudi.pca* function. Broad signals of population genetic structure were investigated using discriminant analyses of principal components (DAPC; Jombart et al 2008). The optimal number of clusters was determined using the *find.cluster* function followed by the *dapc* function using said clusters as the most likely observed structure. The DAPC was visualized using the *scatter* function. A minimum spanning tree calculated from the squared distance between individual was plotted to shed light on the phylogenetic relationship of each inferred cluster. Lastly, the inferred clusters were applied as the population factor and the genetic differentiation between populations (variety names) as well as for the inferred clusters were calculated using the *pairwise.fst* function. Diversity indices for varieties representing putative seed lines for which at least three individuals were available in the dataset were also assessed using the *locus_ table* function, where variety names were used as population indicator.

### Statistical analyses of chemotypic data

A subset of 118 samples from Nevada were also chemotyped at 9 cannabinoid and 17 terpenes, following the methods described by Orser et al., (2018). Since the genetic panel was developed to find the most informative genetic markers associated with chemotypic expression, we grouped individuals according to the clusters from the DAPC and visualized the chemotype variation using side by side boxplots of the top cannabinoid and monoterpenes. Similarly, R was used to read the chemotypic data using the *read.table* function. The *boxplot* function was used to plot the top cannabinoid and terpenes expressed in each cluster.

## RESULTS

### Extent and distribution of genetic diversity and population structure in modern Cannabis

The 22 SNP panel used in this study was selected to represent a broad coverage of the cannabis genome and individual SNPs were found to be located on all cannabis linkage groups with the exception of chromosome 8 (Table 1). As such, levels of polymorphism varied widely between SNPs, from fixed mitochondrial alleles that allow for the discrimination of fibre-type and resin-type cannabis (Figure 1,2,3), to highly variable nuclear markers. Of note, two resin-type landrace varieties from Kyrgyzstan and Egypt were the exception to the rule, both displaying the fibre-type mitochondrial haplotype while expressing THC as the main cannabinoid. Heterozygosity at the nuclear markers ranged from 0.03 to 0.50 (Table 1, Supplementary Table S1). Three markers targeting the THCAS gene cluster offered strong discrimination of major cannabis groups, associated with the two major pentyl cannabinoids THC and CBD. In particular, the *SW6 and VSSL_BtBD* markers were fixed for one allele in all CBD expressing varieties (fibre and resin-types), while being fixed for other allele or heterozygote in all THC expressing varieties. In addition, the SVIP14 locus was also strongly associated with cannabinoid expression data (Table 2).

**Figure 1.**
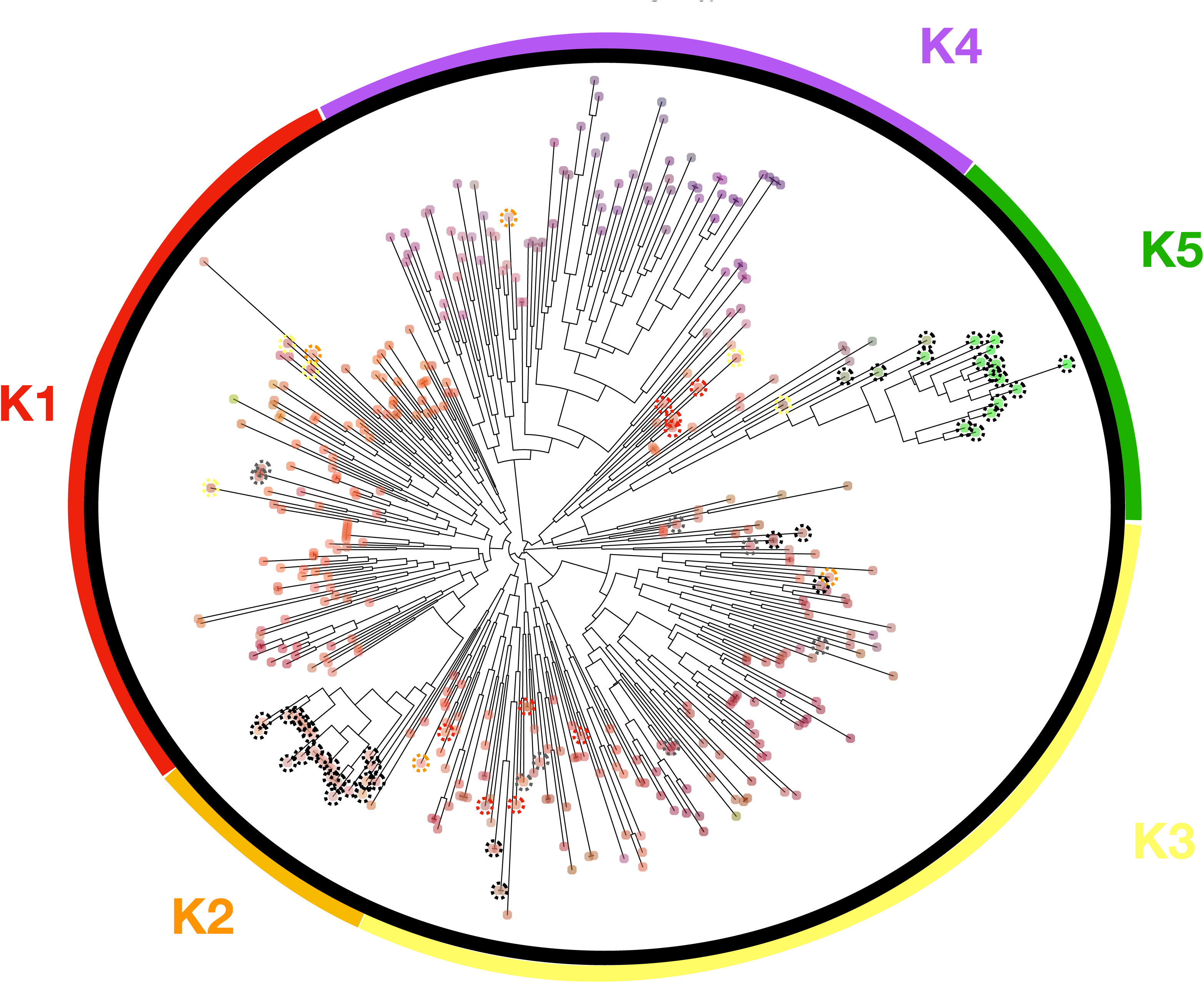
Neighbour-joining tree. Showing the relative location of the 420 Cannabis accessions type at 22SNP. DAPC clusters are shown with K1-K5 represented by different colors. K1-K4 are resin type Cannabis and K5 is the fiber type Cannabis or hemp. Colored dotted circles highlight individuals assigned differently between the neighbor-joining tree and DAPC clusters. Type-III plants are shown with a dotted black circle and type-II plants are shown with dotted grey circle.

**Figure 2.**
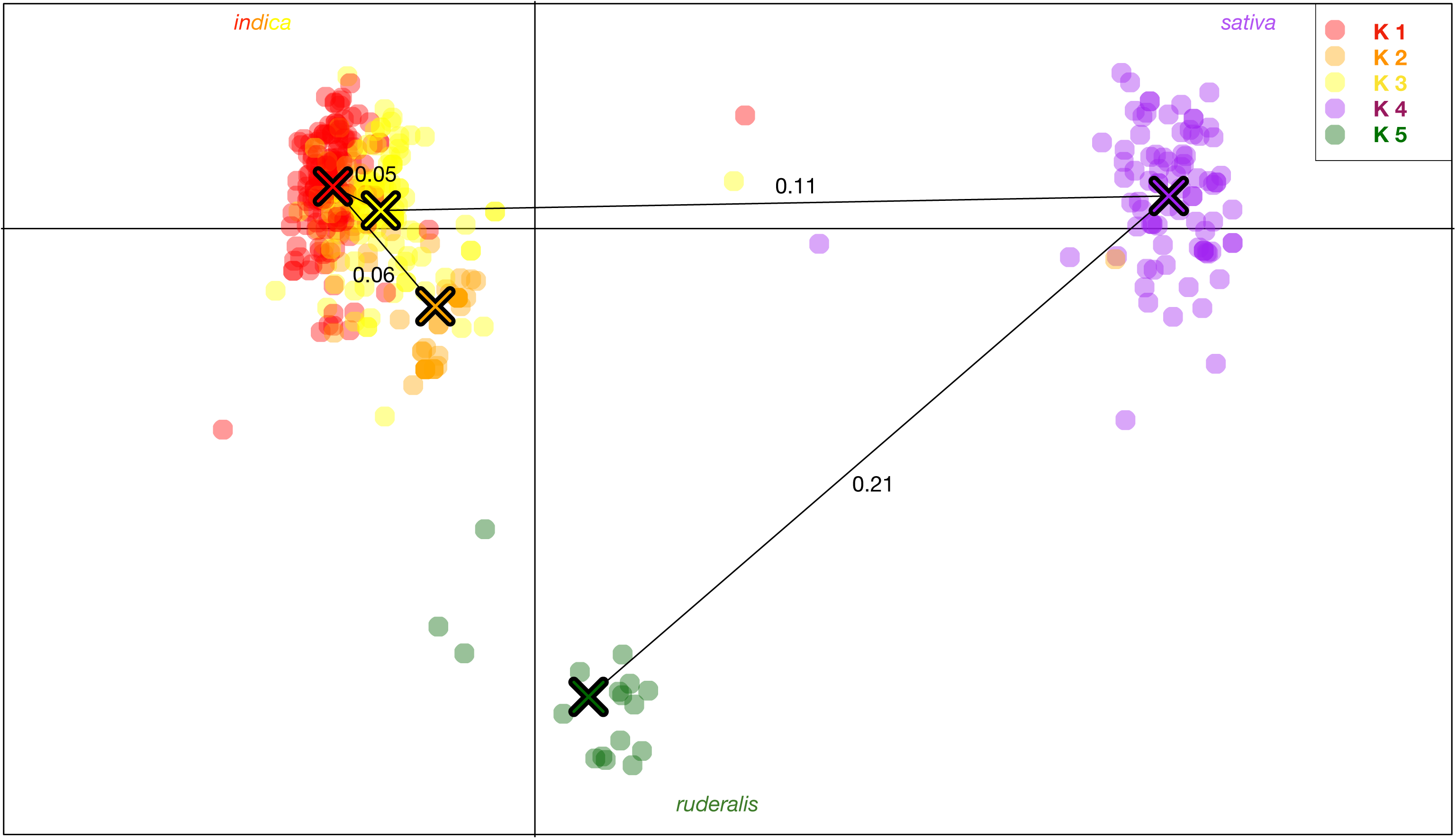
DAPC scatterplot. Showing the relative location of each individual sample in two dimensional space, overlaid by a minimum spanning tree calculated from the squared distance between individual to represent the phylogenetic relationship between inferred clusters. K5, hemp or “*ruderalis*” appears ancestral and the most differentiated group, followed by K4, terpinolene dominant resin accessions. The genetic distance between groups (*Fst*) is indicated on the respective branches of the minimum spanning tree.

**Figure 3.**
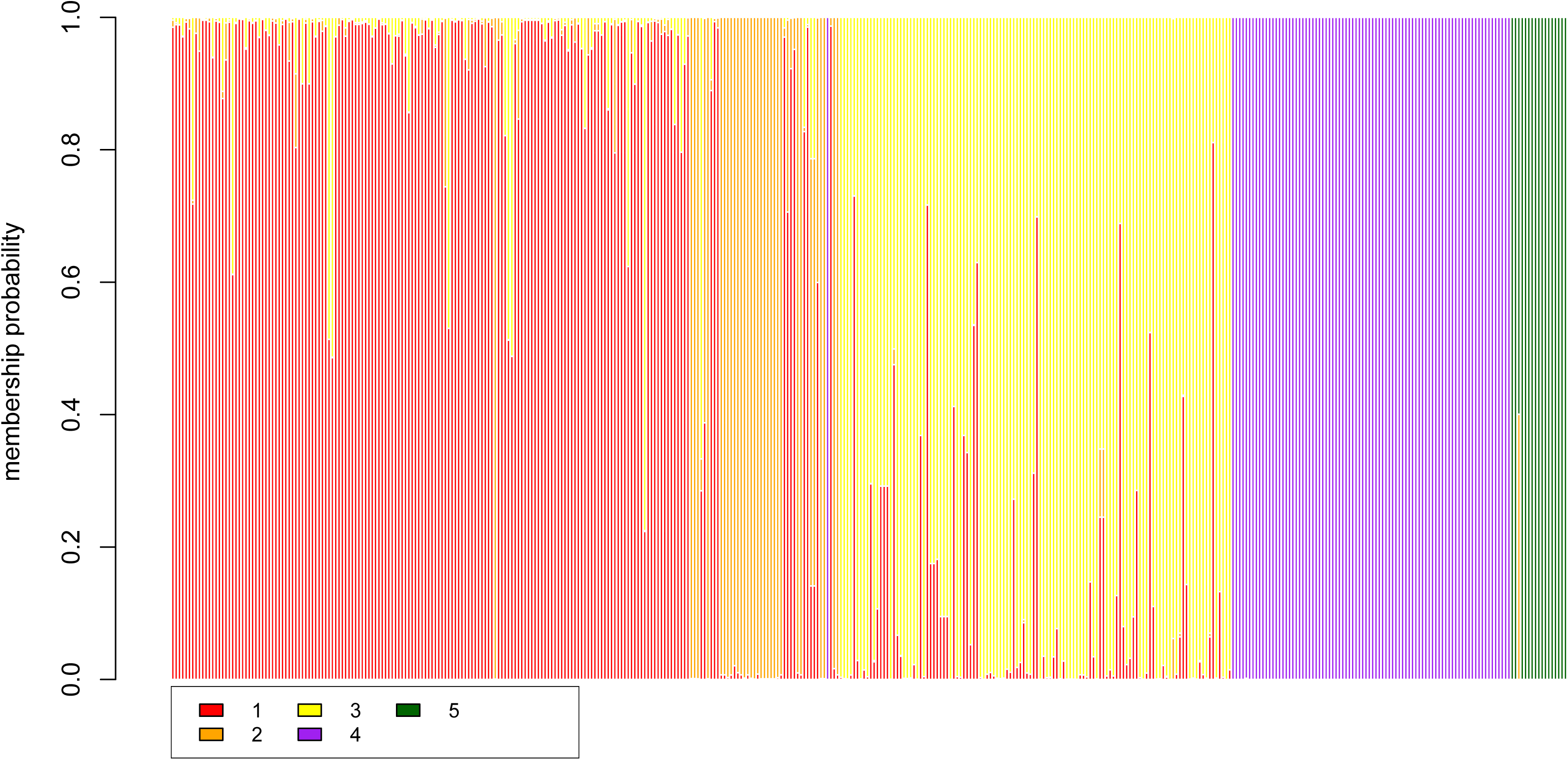
DAPC compoplot. Showing the membership probability of each individual (columns) assignment to each clusters K1 – K5. Mis-assigned individuals can easily be identified as well as F1 hybrids with mixed genotypes.

**Table 1.** Statistics, population genetic metrics and main chemotypes for inferred clusters K1-K5.

| K | N | MLG | H | Hexp | Ia | Terp1 | Terp 2 | Terp 3 | Canna |
| --- | --- | --- | --- | --- | --- | --- | --- | --- | --- |
| 1 | 156 | 135 | 4.8 | 0.31 | 0.22 | Myrcene | Limonene | Linalool | THCA<br>CBCA |
| 2 | 45 | 30 | 3.1 | 0.31 | 1.40 | p-Cymene | Carene |  | CBDA<br>CBCA |
| 3 | 118 | 104 | 4.6 | 0.29 | 0.21 | Myrcene | a-pinene |  | THCA<br>CBCA |
| 4 | 84 | 75 | 4.3 | 0.26 | 0.26 | Terpenolene | Ocimene | Caryophyllene | THCA<br>CBGA |
| 5 | 17 | 17 | 2.8 | 0.133 | 1.12 | Hemp |  |  | CBDA |
| **Total** | 420 | 361 | 5.8 | 0.33 | 0.57 |  |  |  |  |

N - Number of individual samples per population
MLG - the number of multilocus genotypes found in the specified population
H -Shannon-Weiner Diversity index
Hexp-Nei's gene diversity (expected heterozygosity)
Ia - Index of Association for each population factor

**Table 2.** Information about the 22SNPs used in the study. Including genomic location and putative function. Bolded markers indicate those with significant association to the inferred population structure described here.

| Assays ID | SNP | Chromosome | Location | Gene |
| --- | --- | --- | --- | --- |
| SVIP5 | T/A | 5 | 69,688,051 - 69,692,326 | Cannabis sativa uncharacterized LOC115717933 |
| SW6 | G/A | 9 | 25,821,934 - 25,823723 | Inactive THCAS / CBCAS |
| SVIP9 | A/G | 5 | 82,096,056 - 82,098,831 | Cannabis sativa uncharacterized LOC115718065 |
| SVIP10 | C/T | 4 | 10,679,414 - 10,682,584 | Cannabis sativa neurofilament medium polypeptide-like |
| SVIP13 | A/G | 2 | 10,112,681 - 10,121,235 | Cannabis sativa uncharacterized LOC115705170 |
| SVIP14 | A/T | 3 | 417,333 - 420,067 | Cannabis sativa bifunctional endo-1,4-beta-xylanase XylA-like (LOC115710019), mRNA |
| SVIP15 | G/A | 10 | 59,112,921 - 59,117,320 | Cannabis sativa ribose-phosphate pyrophosphokinase 4 |
| SVIP16 | A/C | 10 | 60,829,569 - 60,837,666 | Cannabis sativa probable leucine-rich repeat receptor-like protein kinase At1g35710 |
| SVIP19 | A/G | 1 | 71,846,233 - 71,850,866 | Cannabis sativa mechanosensitive ion channel protein 8-like |
| SVIP21 | G/A | 10 | 58,184,100 - 58,188,483 | Cannabis sativa uncharacterized membrane protein At3g27390 |
| SVIP22 | A/G | 4 | 88,963,964 - 88,967,267 | Cannabis sativa solute carrier family 35 member F5 |
| SVIP23 | C/T | 10 | 47,593,480 - 47,598,098 | Cannabis sativa Cs2S genes for albumin |
| VSSL_BtBd | C/T | 9 | 25,821,934 - 25,823723 | Inactive THCAS / CBCAS |
| VSSL_A250D | C/A | 9 | 25,821,934 - 25,823723 | Inactive THCAS / CBCAS |
| VSSL_mito | C/A | Mitochondria | 317,914 - 318,214 | Downstream of trnC tRNA |
| VSSL_digi2 | C/A | 5 | 14,237,657 - 14,252,007 | Cannabis sativa O-glucosyltransferase rumi homolog |
| VSSL_digi3 | T/C | 6 | 27,445,636 - 27,447,293 | Cannabis sativa uncharacterized LOC115719990 |
| VSSL_digi4 | T/A | 10 | 56,459,661 - 56,460,726 | Cannabis sativa uncharacterized LOC115700304 |
| VSSL_digi6 | C/T | 7 | 1,868,696 - 1,880,067 | Cannabis sativa transcriptional corepressor LEUNIG_HOMOLOG |
| VSSL_digi7 | G/A | 6 | 74,036,351 - 74,039,762 | Cannabis sativa pentatricopeptide repeat-containing protein At5g59600 (LOC115718943), transcript variant X2 |
| VSSL_digi12 | T/C | 5 | 37,063,921 - 37,071,583 | Cannabis sativa K(+) efflux antiporter 5-like |
| VSSL_digi14 | C/T | 3 | 211,984 - 216,544 | Cannabis sativa putative disease resistance RPP13-like protein 1 |
| VSSL_digi19 | G/A | 7 | 56,524,106 - 56,525,416 | Cannabis sativa uncharacterized LOC115722935 |

The DAPC exercise clustered cannabis varieties into five groups (Figure 1,2,3), which was mostly congruent with the independent neighbor joining tree (Figure 1). European Hemp (K5; 15 individuals, C. s. *ruderalis*, typically fibre or grain cultivars, often autoflowering) was clearly distinct from all drug-type cannabis accessions, including high CBD resin expressing accessions. Interestingly resin (drug)-type cannabis consisted of four main genetic clusters, K1 and 3 (156, 118 individuals,, (myrcene/limonene/linalool and myrcene/pinene dominant respectively) which can be considered having a *C. s. indica* phenotype and perceived effect, while K4 (84 individuals, terpinolene) contain mainly accessions of equatorial or *C. s. sativa* designation and phenotype as well as hybrids. K2 (45 individuals, cymene dominant) consisted mostly of the high CBD resin “hemp” from the United States (Figure 4). One known first generation hybrid (“S2”) between an autoflowering male “*Darryl*” and a CBD resin type named “*Intergallactic Princess*” (not sampled here) was found to be assigned to both K2 and K5 in a 40:60 proportion skewed towards the father’s origin (Figure 3). Other possible F1 hybrids were detected between K1 and K3 as well as possibly mis-assigned THC resin individuals into the K2 cluster (Figure 1,2,3).

**Figure 4.**
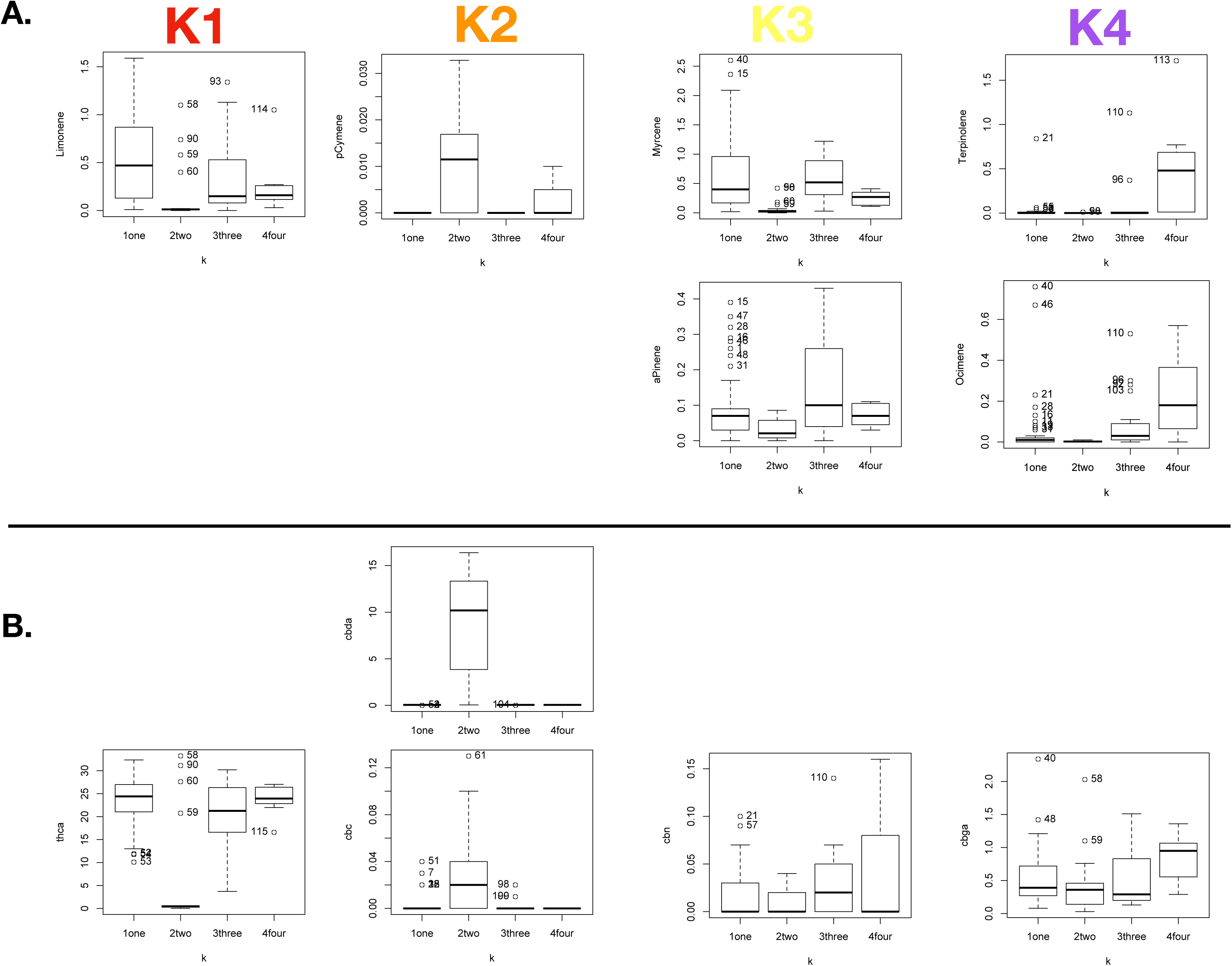
Boxplots of chemotypic data for each inferred K1-K4. No chemotype data was available for K5, yet all individuals from that cluster are expected to display a low resin type-III phenotype. A. Total terpene percentage per dry weight content as determined by GC-MS. B. Total cannabinoid percentage per dry weight content as determined by HPLC.

### Multilocus genotypes, identification of identical clones

In total, 361 multilocus genotypes (putative clonal lines) were identified in the 420 Cannabis samples. While fourteen pairs of known labelled clones were confirmed using the 22 SNP assay, mislabelled accessions with identical multilocus genotypes were frequently detected as follows: “*Unidentified*” and “*Hindu Kush*”, “*GGC*” and “*Purple God*”, “*Atomical Haze*” and “*Tangerine Dream*” and “SFVOG“, “*Gorilla Glue*” and “*Holy Grail*”, “*Agent Orange*” and “*Girl Scout Cookies*”, “*UK Cheese*” and “*Jamaican Ten Speed*”, “*Chem 91”* and “*Colorado Sunset*”, “*Jet Fuel*” *and* “*Louis VIII*”, “*Blackberry Cream*” and “*Slime Dawg MillaNaire*”, “*Tangerine Dream*” and “*Violator Kush*”, “*Original Amnesia*” and “*Sour Tangie*”, “*Billy Crystal*” and “*Blueberry Kush*”, “*5^th^ Dimension*” *and* “*Gorilla Glue*”, “*Garlic*” *and* “*Gelato Breath*”, “*Blue Dream*” with two “*Blue Hash Plant*” samples, seven samples including five labelled “*Pink Kush*”, one mislabelled “*Atomical Haze*” and one *“LA Lights”,* seven unlabelled Resin Hemp from Nevada, seven unlabelled resin Resin hemp samples including one labeled *“Cherry Wine”,* as well as three Resin Hemp samples labelled “Alamo”, “Adam” and “Shore”.

### Diversity within seed lines and inferred clusters

Twenty of the 22 markers were found to deviate from Hardy-Weinberg equilibrium (HWE) in at least one of the 71 populations/seed lines (Supplementary Figure S1), which was not surprising in itself, given the domestication history and strong selective forces for chemotypic expression in modern North American commercial Cannabis cultivars. Of interest when repeated in the larger clusters determined using DAPC, a total of four markers were found to not deviate from HWE (Supplementary Figure S2). The average heterozygosity within seed lines (putative populations) was 0.33, which was considered much higher than what was to be expected in any other major stable commercial crops. Interestingly, the most homozygous line, with heterozygosity of 0.09 was the Canadian fiber/grain cultivar “*X59*” (Supplementary Material Table S1, Table S2). Several drug cultivars, including “*Pink Kush*”, “*Punch Breath*”, “*Durga Matta II CBD*”, “*Durga Matta*”, “*Cotton Candy*”, “*Chem4OG*”, “*33^rd^ Degree”* and “*ASD*” all from known seed banks displayed relative good stability with heterozygosities below 0.2. Another metric of interest is the index of association (*Ia*; Brown, 1980). This index brings an additional insight as a tool to quantify the reshuffling of alleles that occurs in sexually outcrossing species. A deviation from zero (typical of clonal population) indicates increased genetic distance between two individuals from the same seed line. Once again “*X59*” displayed the least distance between individuals indicating a possible strong selection for stable traits in the cannabinoid, fiber and grain expression pathways, and thus a good homogeneous production. For drug-type varieties, the three “*Durga Matta II CBD*” accessions, which were vegetative cuttings from the same mother plants were as expected confirmed to be identical clones. On the other end of the spectrum, several drug-type cultivars had very large *Ia*, which may indicate mislabelling of individual plants or tremendous outcrossing, a syndrome of using F1 hybrids, which appears quite common in the industry to date.

### Association between genetic clusters and chemotypic expression

Looking through a broader lens at the 5 clusters into which the 420 samples segregate one can clearly see a strong differentiation between fiber and resin type Cannabis (Figure 1,2,3, Table 1). One can infer strong selective pressure against THCA expression in K2 (CBD resin type) and K5 (Industrial hemp). Individuals in these clusters, while expressing similar chemotypes, likely underwent a bottleneck for CBDA expression, while displaying large *Ia* values, likely indicative of the polyphyletic and broad origins of the samples at hand for both the resin and fiber type cannabis. While no chemotypic data was available for the fiber type cultivars from K5, a subsample of 118 resin type cultivars with chemotypic data, particularly for major cannabinoid and terpenoid expression demonstrate that (K2 CBD resin type) also consistently expressed p-cymene more so than other resin type accessions (Figure 4, Table 1). Among the THC expressing resin type cluster, K4, the terpinolene dominant group also appeared to accumulate more CBGA and less CBCA than K1-3 (Figure 4, Table 1).

## DISCUSSION

The Cannabis (2n = 2x = 20) draft genome has a haploid genomic sequence of over 876Mb – 1000Mb (Laverty et al., 2019; McKernan et al., 2020) and transcriptome of at least 30,000 genes (van Bakel et al., 2011, Jenkins and Orsburn, 2019a,b,c). The genome displays large amount of polymorphism with a single nucleotide polymorphism (SNP) present every one in a hundred to one in fifty base pairs (McKernan et al., 2020). The phylogenetic relationship and basis for the infra-genus classification have typically recognized a broad structure with divergence between fiber type hemp and drug/resin types Cannabis (Sawler et al., 2015; Dufresnes et al., 2017). In the present study, we delve deeper into the extent and distribution of genetic diversity in modern commercial Cannabis using a novel targeted genetic assay.

While often debated in the literature and confused by lore, our data supports a strong historical and genome-wide division between fiber and resin type cannabis. The maternally inherited mitochondrial DNA supports the ascertion of McPartland and colleagues (2018) which suggests that hemp (*C. s. ruderalis*) is the ancestral group and originated in Europe about 19.7M years ago. A combination of genetic drift and selection then likely contributed to the observed differentiation between fiber and resin cultivars (McPartland et al., 2018). The introgression of an active CBDAS into resin type cannabis likely occurred over the past decade since the advent of medical and recreational Cannabis legislation in Europe and North America. Of interest high CBD and balanced (Type II) accessions were found to cluster into the three resin groups identified here, suggesting a polyphyletic origin of high CBD resin type Cannabis. It is assumed from mapping population that the active form of CBDAS and THCAS are at different loci on Chromosome 9, 8 cM apart in a linked tandem repeat region nestled in a complex array of transposable elements (Weiblen et al., 2015), making the characterization of this region quite complex. Yet, further whole genome sequencing data, particularly using long reads has enabled deeper insight into the structure of the cannabinoid cassette, and demonstrates that the inactive CBDAS gene is in close linkage to the active THCAS (McKernan et al., 2020).

In addition to cannabinoid expression, another marker linked to xylan polysaccharide metabolism (SVIP14; 1-4 Beta Xylanase) was found to contribute to the separation between resin and fiber types which may play a role in fiber quality, given its putative function of breaking down the major constituent of cell walls. Such marker may provide a possible avenue for the development of multi-purpose resin/fiber cultivars.

Integrative analyses revealed a co-expression network of genes involved in the biosynthesis of both cannabinoids and terpenoids from common precursors (Zager et al., 2019). As such, we searched for signals underlying the within resin type cannabis clustering which can be differentiated by the dominant terpene expression, often under the control of two dozen terpene synthase genes (TPS; Allen et al., 2019). While we did not find specific TPS linked markers, we found that a number of SNPs falling in uncharacterized regions of the current *C. sativa* genome were associated with the differentiation between terpene groups in the resin accessions sampled here. Two markers in particular showed strong differentiation between Terpinolene dominant (*“sativa”*) and the other myrcene and limonene dominant accessions (*“indica”*), in particular VSSL_digi2, located in an O-glucosyltransferase rumi analogue involved in ribosome biogenesis and SVIP16 a protein kinase possibly involved in developmental and defense-related processes.

Additionally, the chemotypic data available in the study supported the assertion by others (McKernan et al., 2020) that the presence/absence of a CBCAS gene in resin type cannabis may be responsible for the “leaky” expression of THCA even in cultivars that do not contain an active copy of THCAS. As such, selection against the presence of the CBCAS may provide a possible avenue towards the development of high resin cultivars that are compliant with the current USDA / Health Canada domestic hemp production programs.

## CONCLUSION

We present a targeted genetic assay and algorithms that inform on the sub-genus classification in Cannabis. We demonstrate the use and repeatability of the assay to tease fiber from resin type cannabis, through their mitochondrial lineages and cannabinoid synthases as well as derive possible chemotype classes within resin type Cannabis. We demonstrate some of the utility of the assay as it related to breeding compliant Cannabis and in providing a rapid means to individually type Cannabis accessions and to derive an individual fingerprint that may be used in seed to sale tracking and traceability endeavours. The population level data demonstrates that most resin type varieties exhibited high heterozygosity and as such should be considered unstable at this stage. The use of our array or similar technologies may help in reducing heterozygosity and improving on the stability of trait expression in a similar manner as has been achieved in a fiber type cultivar sampled here, with low heterozygosity and stable trait expression in large seed batches.

## List of abbreviations

PCR: Polymerase Chain Reaction
SNP: Single Nucleotide Polymorphism
KASP: Kompetitive Allele Specific PCR
DAPC: Discriminant Analysis of Principal Components
PCA: Principal Component Analysis
THC: Tetrahydrocannabinol
CBD: Cannabidiol

## AVAILABILITY OF DATA AND MATERIALS

The terpene dataset for 118 individual samples from Nevada is available at the following can be accessed here (https://doi.org/10.6084/m9.figshare.11780103.v1). The genetic data from the 22 SNPs type in 420 individuals with no missing data can be accessed here (https://doi.org/10.6084/m9.figshare.11778936.v1).

## AUTHOR’S CONTRIBUTIONS

Conceptualization: PH CO. Formal analysis: PH BG AM KH AH SJ DJ. Funding acquisition: PH JB JCRJ IW MKD DG CO. Investigation: PH SK KK BG AM KH AH SJ DJ. Samples and resources: PH DJ JCRJ IW MG JB DC DA MKD MM OWH DG. Writing – original draft: PH. Writing – review & editing: PH SK KK BG AM KH AH SJ DJ JCRJ IW MG JB DC DA MKD MM OWH DG CO.

## ACKNOWLEDGEMENENTS

The authors would like the extend our sincere gratitude to the cultivation partners who contributed samples to the study, in particular the Emerald Flower Farm, Terra Growing, Foreman Farms, Oro Verde, Apogee Life, Kambietz Farms, Flying Creek Trading, Pure Farma Solutions, Good Uncle Green Eyes, Breeder Steve, Matrix NV, Flower One, GLP, Greenway, CCLV, Green and Gold, Nature’s Chemistry, Western State Hemp, Harris Farms, Leafceuticals, Hemp Inc., Calineva Farms, Happy Campers, Yield Farming, Franklin BioScience, Polaris MMJ, Acres, Thompson Farm One, and Green Harvest. Dr. D. Darryl Hudson is thanked for providing a sample of a *ruderalis* male aka “*Darryl*”.

---

**Supplementary Figure S1.**
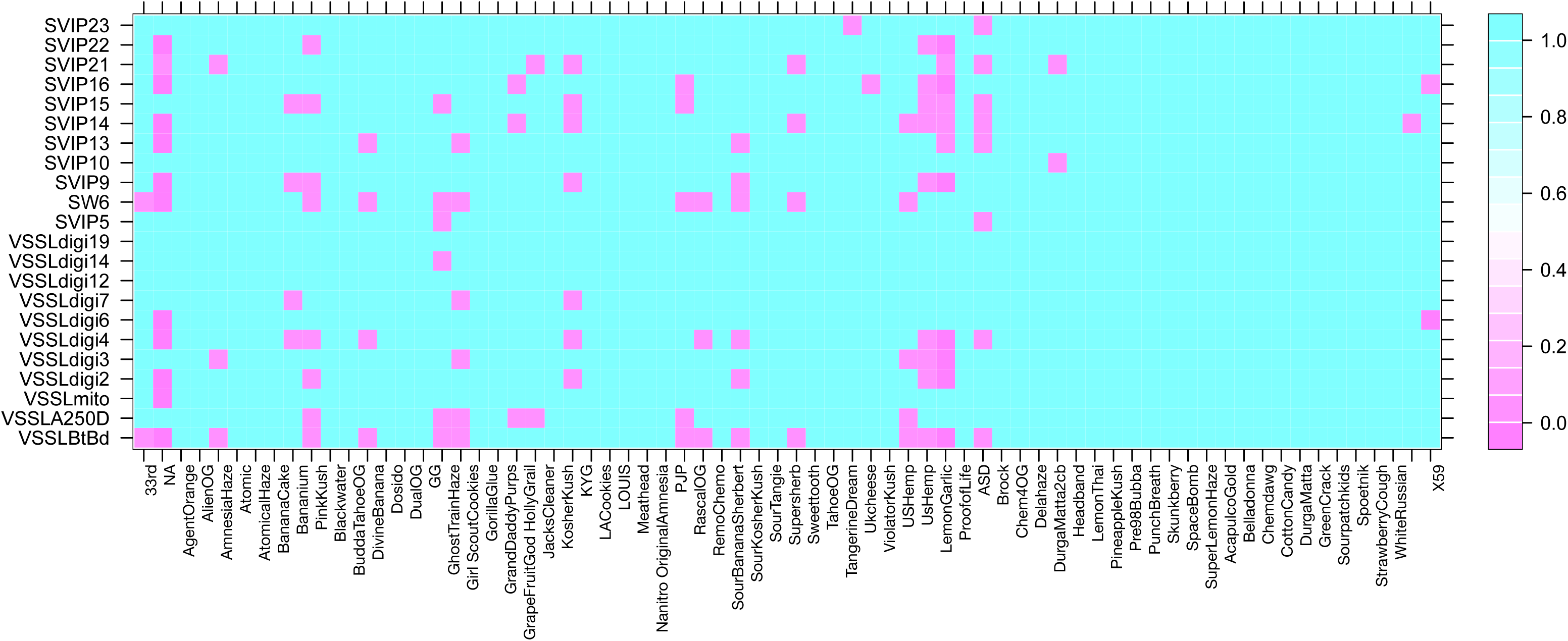
Locus specific deviation from HWE for each samples seed stock. Heat map indicated P-value of test with pink boxes indication significant deviation from HWE.

**Supplementary Figure S2.**
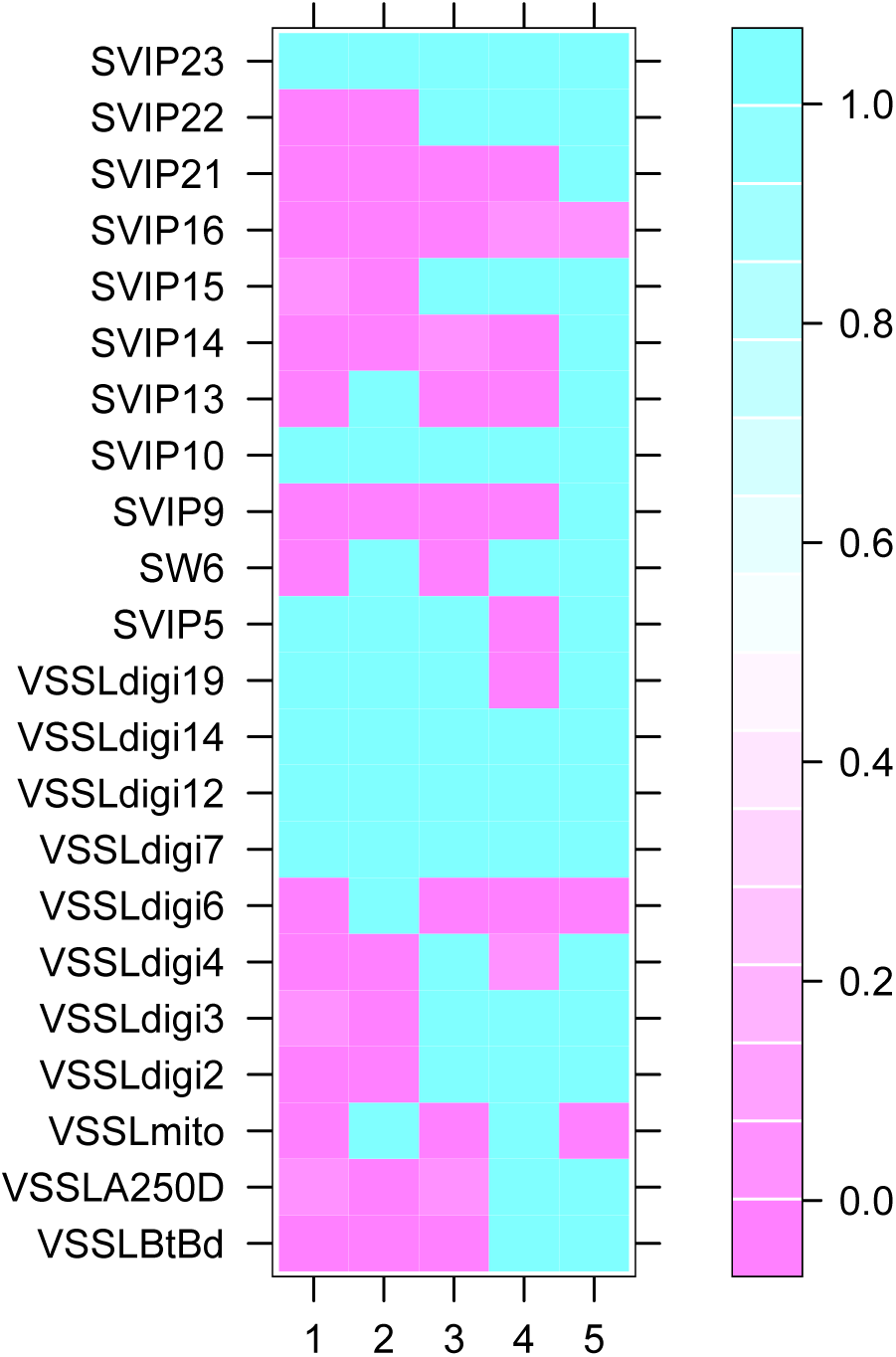
Locus specific deviation from HWE for each inferred clusters. Heat map indicated P-value of test with pink boxes indication significant deviation from HWE.

**Supplementary Table S1.** Seed stock specific population genetic metrics.

| Pop | N | MLG | Hexp | la |
| --- | --- | --- | --- | --- |
| 33rd | 5 | 5 | **0.190909091** | 1.580487805 |
| AmnesiaHaze | 4 | 4 | 0.381493506 | 0.725925926 |
| ASD | 4 | 3 | **0.188311688** | 2.217391304 |
| AtomicalHaze | 3 | 3 | 0.275757576 | 1.818181818 |
| BananaCake | 3 | 3 | 0.284848485 | 0.625 |
| Bananium | 5 | 5 | 0.246464646 | 0.475 |
| Blackwater | 3 | 3 | 0.281818182 | 2 |
| BuddaTahoeOG | 3 | 3 | 0.327272727 | -0.2 |
| Chem4OG | 3 | 3 | **0.163636364** | 2.5 |
| CottonCandy | 3 | 3 | **0.133333333** | 0.75 |
| Delahaze | 5 | 4 | 0.27979798 | 3.058020478 |
| DivineBanana | 4 | 4 | 0.282467532 | 1.402298851 |
| Dosido | 3 | 3 | 0.321212121 | 1.6 |
| DualOG | 6 | 6 | 0.289944904 | 0.8125 |
| DurgaMatta | 3 | 2 | **0.2** | 8 |
| DurgaMatta2cb | 3 | 1 | **0.190909091** | **0** |
| GhostTrainHaze | 4 | 4 | 0.355519481 | 1.663366337 |
| Girl coutCookies | 7 | 7 | 0.308191808 | 1.30044843 |
| GorillaGlue | 3 | 3 | 0.266666667 | 3.454545455 |
| GrapeFruitGod | 4 | 4 | 0.258116883 | 1.454545455 |
| HollyGrail | 4 | 4 | 0.285714286 | -0.295652174 |
| JacksCleaner | 3 | 3 | 0.275757576 | 0.625 |
| KosherKush | 4 | 4 | 0.331168831 | 0.225806452 |
| KYG | 3 | 3 | 0.363636364 | -0.8 |
| LACookies | 5 | 5 | 0.294949495 | -0.032171582 |
| LOUIS | 3 | 3 | 0.272727273 | 8.25 |
| Meathead | 2 | 2 | 0.310606061 | NA |
| Nanitro | 3 | 3 | 0.260606061 | 6.3 |
| OriginalAmnesia | 5 | 4 | 0.214141414 | 1.197452229 |
| PineappleKush | 3 | 3 | 0.3 | 0.5 |
| PinkKush | 7 | 3 | **0.18981019** | 1.5 |
| PJP | 4 | 4 | 0.225649351 | 0.03030303 |
| ProofofLife | 4 | 2 | 0.266233766 | 3 |
| PunchBreath | 3 | 2 | **0.136363636** | 2 |
| RemoChemo | 4 | 4 | 0.251623377 | -0.677419355 |
| SourTangie | 7 | 7 | 0.283716284 | 0.725943971 |
| SpaceBomb | 3 | 3 | 0.233333333 | 1.8 |
| StrawberryCough | 3 | 3 | 0.124242424 | 0.75 |
| Sweettooth | 3 | 3 | 0.351515152 | 1.153846154 |
| TahoeOG | 5 | 5 | 0.211111111 | 1.482758621 |
| TangerineDream | 9 | 9 | 0.322638146 | 0.840565086 |
| UsHemp | 20 | 11 | 0.279545455 | 0.994321219 |
| USHemp | 7 | 6 | 0.293706294 | 1.635103926 |
| ViolatorKush | 7 | 7 | 0.303696304 | 1.38769671 |
| WhiteRussian | 7 | 7 | 0.184315684 | 0.327198364 |
| X59 | 14 | 14 | **0.093073593** | **0.167832168** |
| Total / average | 420 | 361 | 0.330011764 | 0.570375586 |

**Supplementary Table S1.** Locus specific statistics.

Supplementary Material Table S2
| SNP | allele | 1-D | Hexp | Evenness |
| --- | --- | --- | --- | --- |
| VSSLBtBd | 2 | 0.486111111 | 0.486690505 | 0.97290917 |
| VSSLA250D | 2 | 0.318568594 | 0.318948294 | 0.722946759 |
| VSSLmito | 2 | 0.073287982 | 0.073375333 | 0.450170654 |
| VSSLdigi2 | 2 | 0.404036281 | 0.40451785 | 0.836002253 |
| VSSLdigi3 | 2 | 0.497276077 | 0.497868778 | 0.994579203 |
| VSSLdigi4 | 2 | 0.427437642 | 0.427947103 | 0.871561439 |
| VSSLdigi6 | 2 | 0.075484694 | 0.075574664 | 0.453144701 |
| VSSLdigi7 | 2 | 0.25997449 | 0.260284352 | 0.656034447 |
| VSSLdigi12 | 2 | 0.04875 | 0.048808105 | 0.413244143 |
| VSSLdigi14 | 2 | 0.136723356 | 0.136886316 | 0.52560179 |
| VSSLdigi19 | 2 | 0.025847506 | 0.025878313 | 0.367110514 |
| SVIP5 | 2 | 0.499770408 | 0.500366082 | 0.99954101 |
| SW6 | 2 | 0.49744898 | 0.498041887 | 0.994921692 |
| SVIP9 | 2 | 0.470510204 | 0.471071003 | 0.94401974 |
| SVIP10 | 2 | 0.128560091 | 0.128713321 | 0.5166806 |
| SVIP13 | 2 | 0.485306122 | 0.485884556 | 0.971379846 |
| SVIP14 | 2 | 0.355226757 | 0.35565015 | 0.768669616 |
| SVIP15 | 2 | 0.499657029 | 0.500252568 | 0.99931449 |
| SVIP16 | 2 | 0.447573696 | 0.448107157 | 0.90420323 |
| SVIP21 | 2 | 0.409180839 | 0.40966854 | 0.843617072 |
| SVIP22 | 2 | 0.317131519 | 0.317509507 | 0.721223835 |
| SVIP23 | 2 | 0.387752268 | 0.388214428 | 0.812593468 |
| mean | 2 | 0.329618893 | 0.330011764 | 0.760884985 |

allele = Number of observed alleles
Hexp-Nei's gene diversity (expected heterozygosity)
Evenness - Index of Association for each population factor

